# Calculating the divergence time by direct comparison between individuals and the out-group

**DOI:** 10.1101/2021.04.04.438302

**Authors:** Tingting Sun, Qing Liu, Meiqi Shang, Kejian Wang

## Abstract

Investigating the divergence time between different populations has been of fundamental interest in evolution biology. In this study, we reanalysed the mitochondrial and Y chromosome DNA of the 1000 Genomes Project, and revealed that most single nucleotide polymorphisms (SNPs) are specific to minority and would be classified as low-frequency mutations. Using these polymorphisms, we recalculated the divergence time of different populations by direct comparison between individuals and the out-group.

---

Investigating the divergence time between different populations has been of fundamental interest in evolution biology. Single-nucleotide polymorphisms (SNPs) are the most common type of polymorphism and have been widely used to determine the divergence time of different haplogroups in human evolution ^1,2^. To avoid potential errors and accelerate the calculation, low-frequency SNPs that account for less than 1% or 5% of the population are generally discarded^3^. To evaluate the contribution of the low-frequency SNPs in the estimation of divergence time, we reanalysed the single nucleotide variants of 2,534 human individuals released by the 1000 Genomes Project^4,5^. We first analysed the mtDNA of the 2,534 individuals and found a total of 3,586 SNPs. Of these SNPs, 2,913 (81%) were present in fewer than ten individuals (accounting for 0.36% of individuals analysed), and 1,296 (36%) SNPs were even specific to a single individual (Figure 1a). There were 60,505 SNPs found in Y-DNA from 1,233 male individuals. Of these SNPs, 50,667 (84%) were present in fewer than ten individuals (accounting for 0.73% of male individuals analysed), and 33,789 (56%) occurred only in a single individual (Figure 1). These results indicate the majority of the SNPs are specific to minority and would be classified as low-frequency mutations in population genetics.

**Figure 1:**
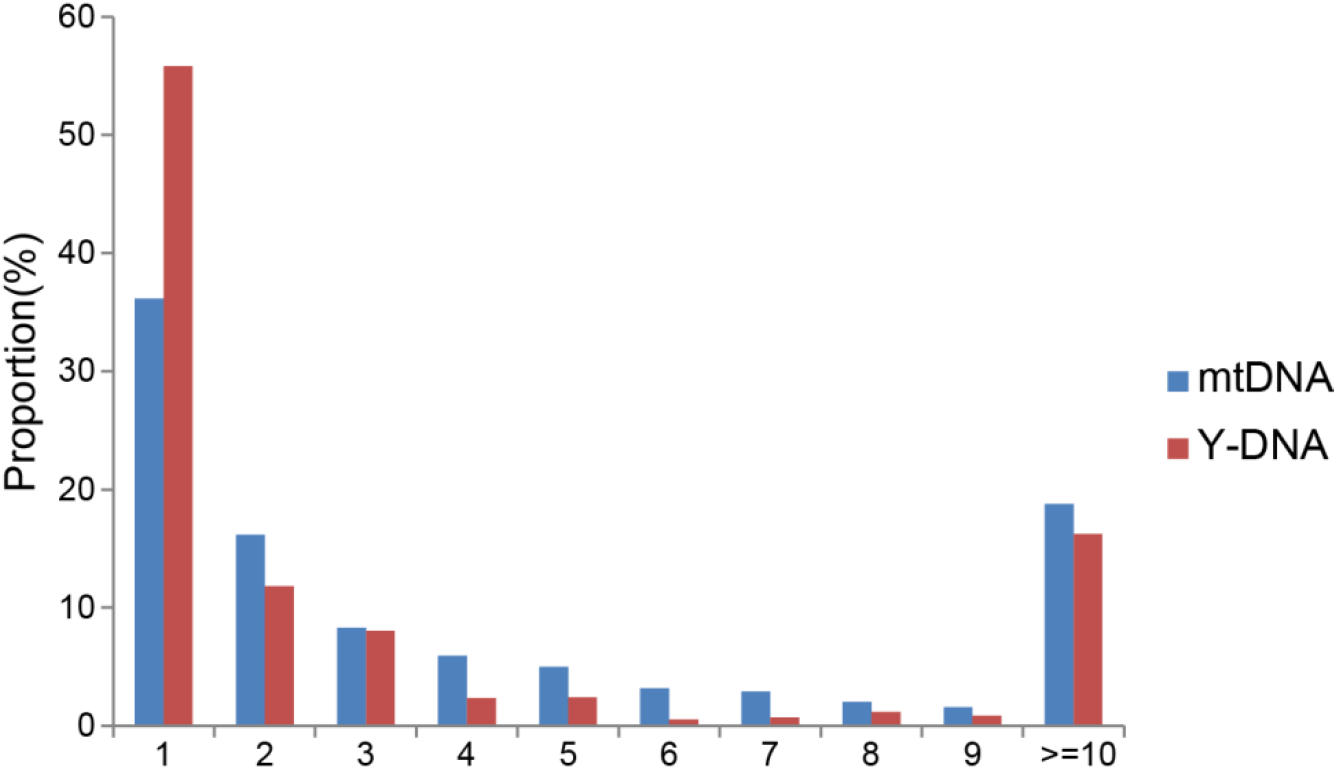
Proportions of specific SNPs in human mtDNA and Y-DNA. The X-axis represents the number of individuals sharing the same SNPs. The Y-axis represents the proportion of SNPs.

Previous studies have shown that the SNPs of the 1000 Genomes Project are supported by high-coverage sequencing data, indicating the accuracy of those SNPs^4^. Here we wanted to further confirm the reliability of those minority-specific SNPs before using them to calculate the divergence time. According to the molecular clock hypothesis, new SNPs are introduced into the genome at a constant rate^6^, it therefore can be inferred that each individual has a similar number of SNP variations compared with the common ancestor. It is also concluded that each individual also shares a similar number of SNP variations with a given common out-group. To verify this, we chose the genome of the chimpanzee as the common out-group because it was the closest living relatives of human beings and had similar mutation rate with human^7^. To prevent any possible errors caused by sequence differences, only SNPs on the same sequence between human and chimpanzee were selected for analysis. In addition, those SNPs located beyond the linkage disequilibrium block were discarded. After screening, a total of 2,854 SNPs of mtDNA and 235,382 SNPs of Y-DNA were remained for further analysis. We compared the SNP differences between human and chimpanzee using those screened SNPs and found that each human individual had a similar number of SNP differences with the chimpanzee, with very limited variation in both mtDNA (921.36 ± 3.79) and in Y-DNA (190,135.15 ± 26.41). This observation indicates that the minority-specific SNPs are reliable, and the mutation rate is similar between different human individuals.

Next, we reanalysed mtDNA and Y-DNA using all screened SNPs and calculated the divergence time of different individuals or groups using the simple model based on molecular clock hypothesis (Figure 2). The principle underlying the model is as follows. If the out-group shares the same mutation rate with the A/B individuals or groups, the number of SNP differences between the out-group (O) and A/B is proportional to twice the divergence time (*T*_1_) between the out-group and A/B. Similarly, the number of SNP differences between A and B is proportional to twice the divergence time (*T*_1_) between A and B. When the split time between the human and the out-group is known, the divergence time between A and B can be determined. The time is calculated according to the following formula:

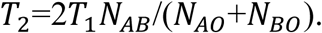

**Figure 2:**
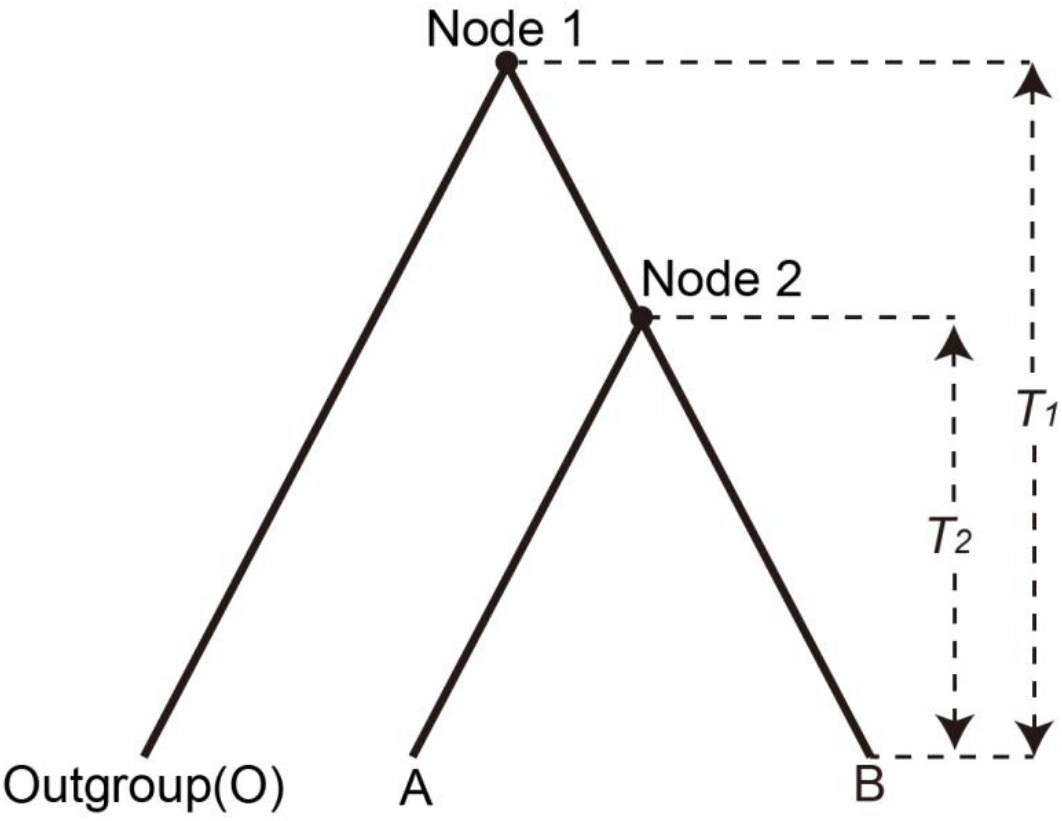
Time calculation model used in this study. A and B represent two different individuals or groups. Node 1 represents the MRCA of Out-group (O) and A/B. The corresponding divergence time is represented as *T*_1_. Node 2 indicates the MRCA of A and B, and the corresponding divergence time is indicated as *T*_1_. If more than two individuals are analysed, the SNP differences between each individual are calculated and then take the average.

To determine the time to common ancestor for modern human, we first constructed a neighbour-joining tree using the mtDNA of all 2,534 individuals and the chimpanzee. The exact time of divergence between humans and chimpanzees remains subject to debate, and the general consensus of five million years was used in this study^8^. The evolutionary tree suggested that the common mtDNA ancestor of modern humans originated in Africa (Figure 3), which was in keeping with previous studies^1,9^. In the polygenetic tree, Node 1 represents the Most Recent Common Ancestor (MRCA) of all human individuals analysed. We calculated the time according to the above formula and found that the MRCA of mtDNA lived about 230 KYA (Figure 3, Table 1 and Supplementary Table S1), which is roughly consistent with previous haplogroup analysis^1,9^. We further constructed neighbour-joining tree using the Y-DNA of 1,233 individuals and the chimpanzee (Figure 4). Using the same calculation, we found that the common patrilineal ancestors of modern humans arose in Africa about 60 KYA (Figure 4, Table 1 and Supplementary Table S1), which is far less than the time (190 KYA) of haploid analysis^4^. The inconsistence might be resulted from the significant variation of the structure and gene content of Y chromosome between chimpanzee and human^10^. It is also likely to be caused by insufficient sample size or sequencing depth of Y-chromosome. Taken together, the direct comparison between individuals and the out-group can be used to estimate the divergence time of mtDNA, but still needs further optimization to calculate that of Y chromosome.

**Figure 3:**
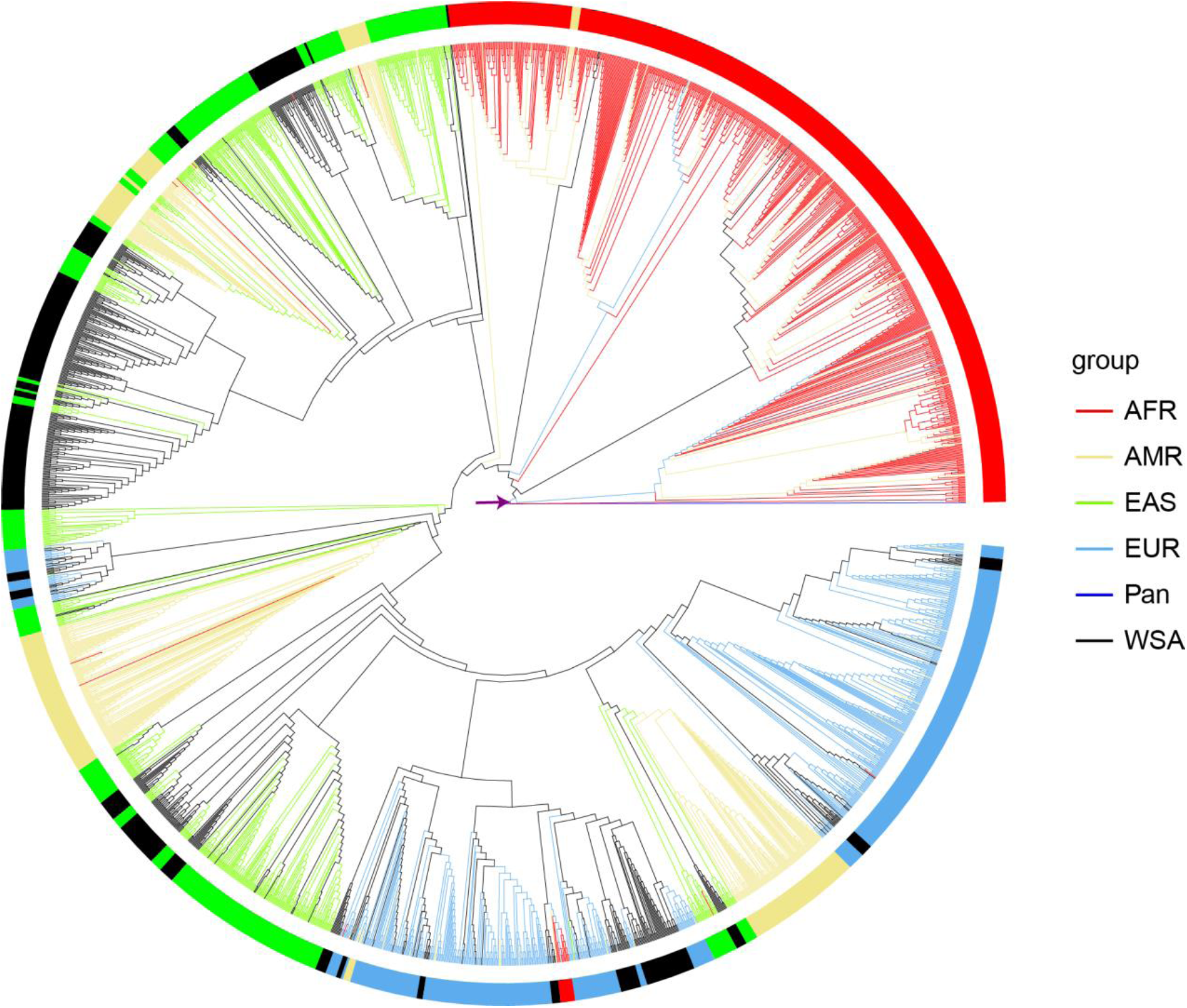
Evolutionary tree based on mtDNA. Populations from Africa (AFR), America (AMR), East Asia (EAS), Europe (EUR), West and South Asia (WSA) were used to construct an evolutionary tree based on mtDNA. The node pointed by the arrow indicates the common ancestor of modern human beings.

**Table 1:**
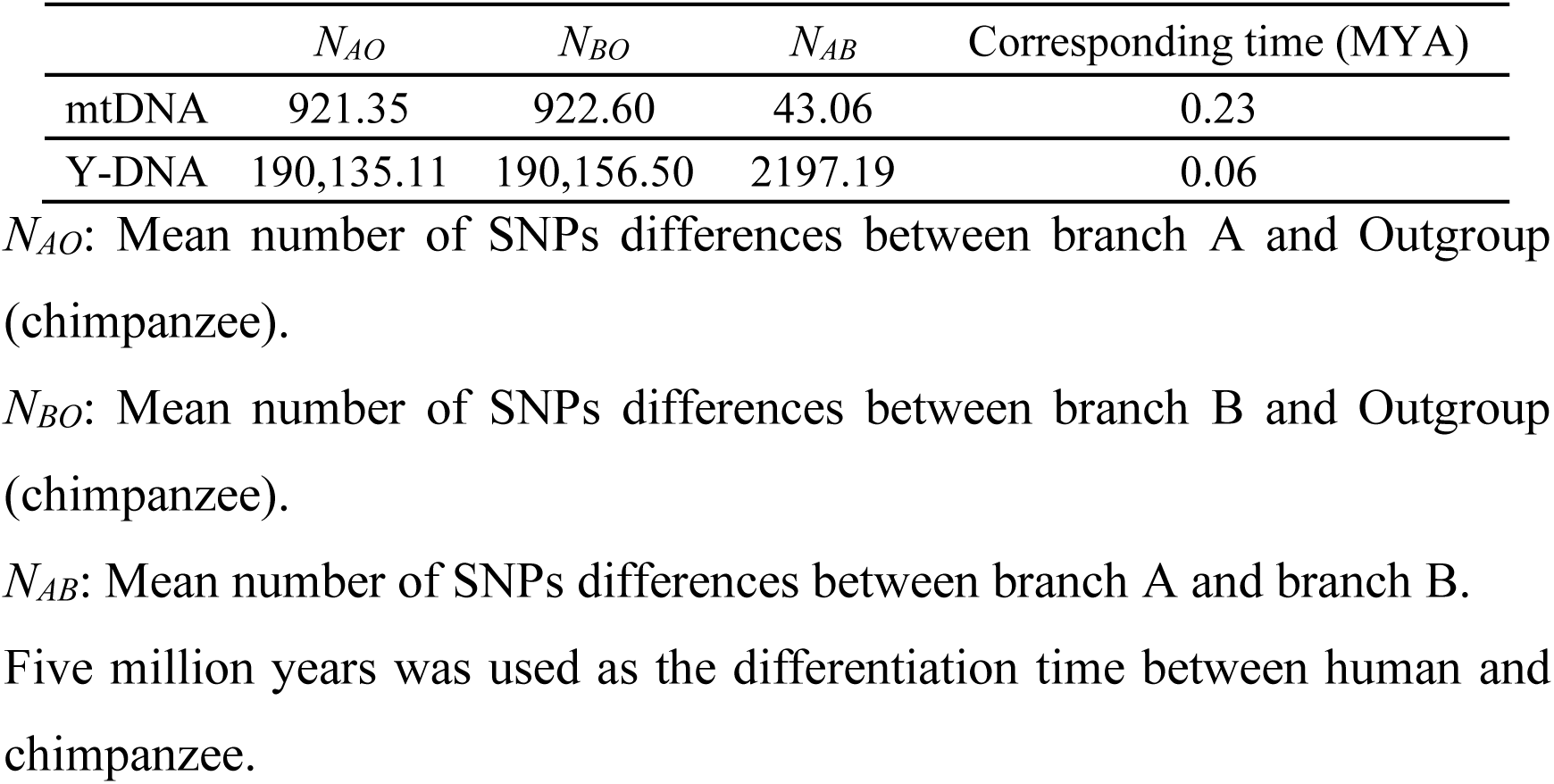
The time to common ancestor for modern human based on evidence from mtDNA and Y-DNA.

| | $N_{AO}$ | $N_{BO}$ | $N_{AB}$ | Corresponding time (MYA) |
| --- | --- | --- | --- | --- |
| mtDNA | 921.35 | 922.60 | 43.06 | 0.23 |
| Y-DNA | 190,135.11 | 190,156.50 | 2197.19 | 0.06 |

$N_{AO}$ : Mean number of SNPs differences between branch A and Outgroup (chimpanzee).
$N_{BO}$ : Mean number of SNPs differences between branch B and Outgroup (chimpanzee).
$N_{AB}$ : Mean number of SNPs differences between branch A and branch B.
Five million years was used as the differentiation time between human and chimpanzee.

**Figure 4:**
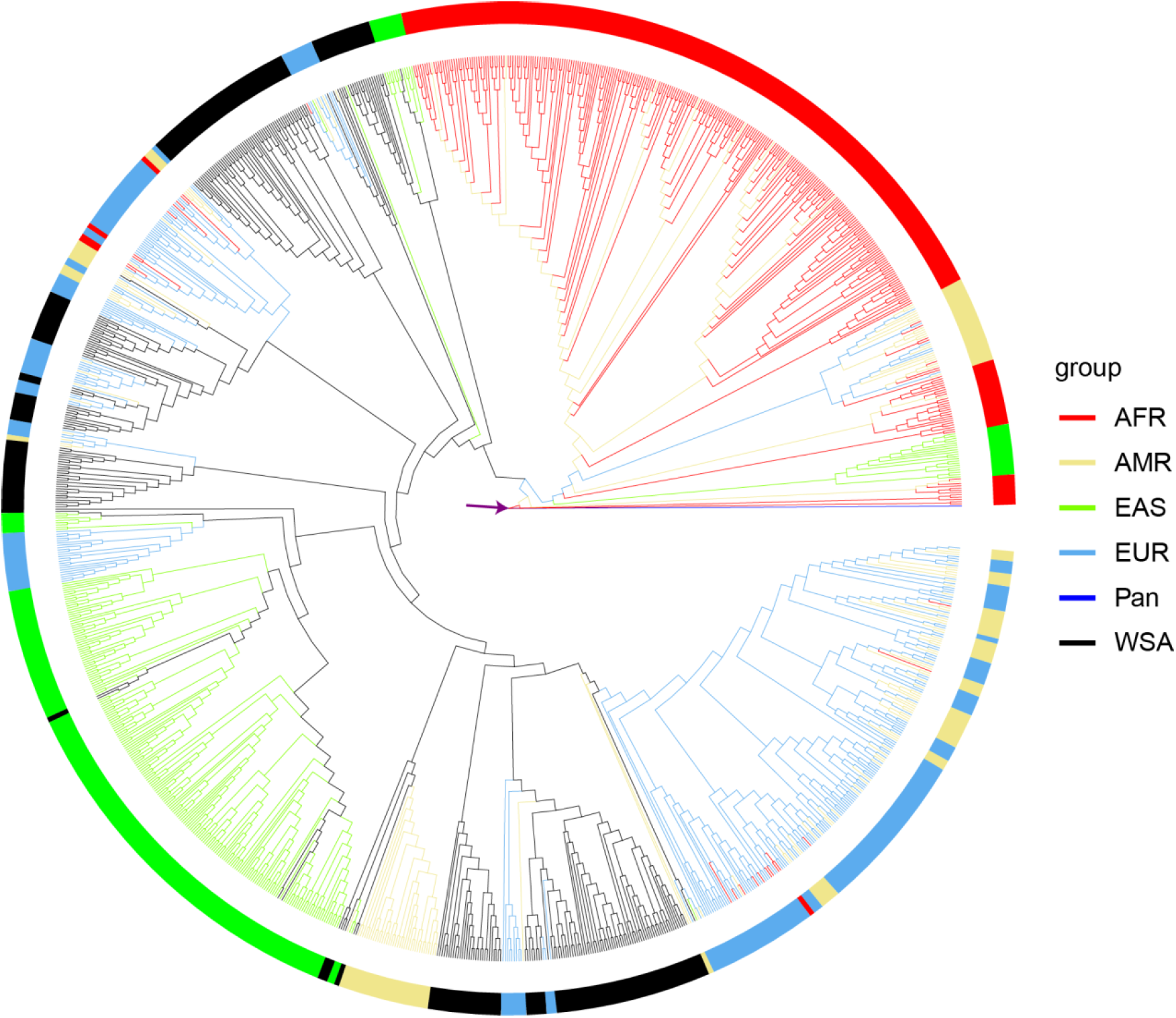
Evolutionary tree based on Y-DNA. Populations from Africa (AFR), America (AMR), East Asia (EAS), Europe (EUR), West and South Asia (WSA) were used to construct an evolutionary tree based on Y-DNA. The node pointed by the arrow indicates the common ancestor of modern human beings.

## Methods

### Read mapping and SNP calling for out-group

The sequencing of 24 chromosomes and the mitochondrial genome—chromosomes 2A and 2B merged into chromosome 2—of the chimpanzee (GCF_002880755.1_Clint_PTRv2_genomic.fna https://ftp.ncbi.nlm.nih.gov/genomes/all/GCF/002/880/755/GCF_002880755.1_Clint_PTRv2/GCF_002880755.1_Clint_PTRv2_genomic.fna.gz) was simulated using ART software with the parameters ‘–l 150’, ‘–f 60’, ‘–m 500’, ‘–s 10’, ‘-p’, ‘-na’ and ‘–ef’^11^. The high quality paired-end reads were aligned to the hs37d5 reference genome using BWA (v0.7.17) with the parameters: ‘mem -t 16 -k 32–M’^12^. PCR duplicates were removed using Picard tools (http://broadinstitute.github.io/picard/). We performed SNP calling using a HaplotypeCaller approach as implemented in the package GATK^13^. A Perl programme was used to filter potentially false SNPs using the following criteria: a homozygous genotype; supporting reads for the reference or alternative allele greater than 30; and the ratio of them was greater than 3.

### Population genetic analysis

Single-nucleotide polymorphism genotyping data available from the HapMap3 Project (Altshuler et al. 2010) was downloaded from the NCBI (ftp://ftp.1000genomes.ebi.ac.uk/vol1/ftp/release/20130502/)^5^. The linkage disequilibrium (LD *r*^2^) of all SNPs in the mitochondrial genome within 50,000 kb were calculated using the Haploview software^14^. For the Y chromosome, 200 SNPs were randomly selected from the SNP genotyping dataset to calculate the LD within 50,000 kb using Haploview. The final SNP dataset was formed by using SNP located in the homologous sequences between human and chimpanzee in the LD block. The IBS distance matrix of the mitochondrial genome and the Y-DNA was computed using PLINK on the SNP dataset which was located in the LD block^15^. The same distance matrix was used to construct a phylogenetic tree by the neighbour-joining method, implementing in PHYLIP and visualised in R^16,17^.

### Analysis of SNP differences between different groups

Individual-specific SNPs and the number of SNP differences between each individual and chimpanzee were calculated using a Perl script. When counting the number of SNP differences, genotypes designated ‘-’, were not counted. For branches with more than one individual, the average number of SNP differences over all individuals was calculated.

## Supporting information

Individuals of two branches from each differentiation nodes of mtDNA and Y-DNA

## Author Contributions

TS conceived the study, conducted the analysis and wrote the paper; QL and MS discussed the results and verified the data; KW conceived and supervised the study; All authors reviewed and edited the manuscript.

## Additional files

Supplementary Table S1: Individuals of two branches from each differentiation node of mtDNA and Y-DNA

